# Altered Temporal Variability of Local and Large-scale Resting-state Brain Functional Connectivity Patterns in Schizophrenia and Bipolar Disorder

**DOI:** 10.1101/2020.02.04.934638

**Authors:** Yicheng Long, Zhening Liu, Calais Kin-yuen Chan, Guowei Wu, Zhimin Xue, Yunzhi Pan, Xudong Chen, Xiaojun Huang, Weidan Pu

**Affiliations:** Department of Psychiatry, The Second Xiangya Hospital, Central South University, Changsha, Hunan, China; Mental Health Institute of Central South University, Changsha, Hunan, China; Department of Psychology, The University of Hong Kong, Hong Kong, China; Medical Psychological Center, The Second Xiangya Hospital, Central South University, Changsha, China

**Keywords:** dynamic functional connectivity, schizophrenia, bipolar disorder, thalamus, sensorimotor, basal ganglia

## Abstract

Schizophrenia and bipolar disorder share some common clinical features and are both characterized by aberrant resting-state functional connectivity (FC). However, little is known about the common and specific aberrant features of the dynamic FC patterns in these two disorders. In this study, we explored the differences in dynamic FC among schizophrenia patients (*n* = 66), type I bipolar disorder patients (*n* = 53) and healthy controls (*n* = 66), by comparing temporal variabilities of FC patterns involved in specific brain regions and large-scale brain networks. Compared with healthy controls, both patient groups showed significantly increased regional FC variabilities in subcortical areas including the thalamus and basal ganglia, as well as increased inter-network FC variability between the thalamus and sensorimotor areas. Specifically, more widespread changes were found in the schizophrenia group, involving increased FC variabilities in sensorimotor, visual, attention, limbic and subcortical areas at both regional and network levels, as well as decreased regional FC variabilities in the default-mode areas. The observed alterations shared by schizophrenia and bipolar disorder may help to explain their overlapped clinical features; meanwhile, the schizophrenia-specific abnormalities in a wider range may support that schizophrenia is associated with more severe functional brain deficits than bipolar disorder.

## 1. Introduction

Schizophrenia and bipolar disorder are two of the most disabling psychiatric disorders worldwide, which are often misdiagnosed in clinical practice because of their overlap in clinical features. These common features entail both cognitive deficits and psychotic symptoms including hallucinations, delusions and disorganized thinking (1–3). Over the years, neuroimaging studies using resting-state functional magnetic resonance imaging (rs-fMRI) have provided evidence for both shared and distinct disturbances in brain functions, as characterized by aberrant resting-state functional connectivity (FC), in the schizophrenia and bipolar disorder (4–7). For instance, when compared with healthy subjects, over-connectivity between the thalamus and sensorimotor cortices was commonly found in both schizophrenia and bipolar disorder patients (4). On the other hand, other unique abnormalities such as hypo-connectivity within frontal–parietal areas were shown only in schizophrenia but not bipolar disorder patients (7). Appreciably, these findings have significantly advanced our understanding of the complex relationship between these severe disorders.

Most previous rs-fMRI studies were performed under the assumption that patterns of brain FC are stationary during the entire scanning period. Yet, it has been newly proven that the brain FC fluctuates over time even during the resting-sate, implying that conventional static FC methodology may be unable to fully depict the functional architecture of brain (8,9). Therefore, the “dynamic FC” has become a hot-spot in rs-fMRI studies to capture the temporal fluctuations of brain FC patterns during the scan (10). Notably, the dynamic features of FC have been associated with a wide range of cognitive and affective processes such as learning (11), executive cognition (12), psychological resilience (13) and emotion (14), as well as multiple common psychiatric and neurological disorders such as autism (15), Alzheimer’s disease (16) and major depressive disorder (17,18). These findings thus highlight the importance of studying dynamic FC for further improving our understanding of both brain functions and dysfunctions.

Despite the accumulating knowledge on dynamic FC, it remains little known about if there are common and/or specific changes in dynamic features of FC in schizophrenia and bipolar disorder. To our knowledge, there have been only a limited number of efforts to date to differentiate schizophrenia and bipolar disorder by features of dynamic FC (19–21). Furthermore, all these studies mainly focus on the dynamic “connectivity state” changes based on the whole-brain FC profiles; therefore, although features of such global connectivity states have been reported to provide a high predictive accuracy in classifying schizophrenia and bipolar disorder (19–21), how these two disorders differ from each other in terms of dynamic connectivity profiles within particular brain regions or systems remains poorly understood, and needs to be further investigated.

The above concerns can be addressed by a novel approach, as proposed in some latest works (22,23), to investigate dynamic FC by defining and comparing the temporal variability of FC patterns involved in specific brain regions or large-scale brain networks. This approach allows localization of those brain regions or networks showing significant group differences in FC variability, thus being helpful to identify aberrant dynamic FC patterns from the perspectives of both local and large-scale brain functional dynamics (23). In fact, using such an approach, the patients with schizophrenia have been recently found to show increased FC variabilities in sensory and perceptual systems (e.g. the sensorimotor network and thalamus) and decreased FC variabilities in high-order networks (e.g. the default-mode network) than healthy subjects at both regional and network levels (22). But to our knowledge, it remains unclear and needs to be tested whether these dynamic changes would be specific to schizophrenia, or shared with bipolar disorder.

Therefore, in this study, we aimed to explore the common and specific dynamic features of both local and large-scale resting-state FC, in terms of temporal variability, the schizophrenia and bipolar disorder. To reach this goal, groups of schizophrenia patients, bipolar disorder patients and healthy controls were recruited and scanned using rs-fMRI; applying a recently proposed novel methodological approach (22,23), temporal variabilities of FC patterns were then compared among the groups at all the regional, intra-network, and inter-network levels. It was anticipated that our results would provide important complementary information to prior studies that mainly focused on the global dynamic FC states (19–21), and further improve our understanding about the relationship between schizophrenia and bipolar disorder from a dynamic brain functional perspective.

## 2. Materials and Methods

### 2.1. Subjects and measurements

According to the Diagnostic and Statistical Manual of Mental Disorders-IV (DSM-IV) criteria, 78 patients with schizophrenia and 60 patients with type I bipolar disorder were recruited from the Second Xiangya Hospital of Central South University, Changsha, China; 69 age-, sex- and education-matched healthy controls without any family history of psychiatric disorders were also recruited from the Changsha city. All participants were right-handed, Han Chinese adults with at least 9 years of education. All participants had no history of any substance abuse, any other neurological disorder, any contraindication to fMRI scanning or any history of electroconvulsive therapy. Because of excessive head motion (see section 2.2), 12 schizophrenia patients, 7 bipolar disorder patients and 3 healthy controls were excluded, and the final analyzed sample consisted of 66 schizophrenia patients, 53 bipolar disorder patients and 66 healthy controls.

For the schizophrenia patients, severity of the current clinical symptoms was assessed using the Scale for Assessment of Positive Symptoms (SAPS) and the Scale for Assessment of Negative Symptoms (SANS) (24). For the patients with bipolar disorder, severity of the current mood and mania symptoms was assessed using the 17-item Hamilton Rating Scale for Depression (HAMD) (25) and the Young Mania Rating Scale (YMRS) (26), respectively. Dosages of antipsychotics in all patients were converted to chlorpromazine equivalence (27). In addition, all participants completed the Information (WAIS-I) and Digit Symbol (WAIS-DS) subtests of the Wechsler Adult Intelligence Scale (28), which measure two important domains of cognitive functions, verbal comprehension and processing speed, respectively (29,30).

The study was approved by the Ethics Committee of the Second Xiangya Hospital of Central South University, and written informed consent was obtained from all participants.

### 2.2. Data acquisition and preprocessing

The details about brain imaging data acquisition and preprocessing can be found in one of our recently published studies (13). Briefly, rs-fMRI and T1-weighted structural images were scanned for each participant on a 3.0 T Philips MRI scanner (repetition time = 2000 ms, echo time =30 ms, slice number =36, field of view =240×240 mm^2^, acquisition matrix =144×144, flip angle =90°, and number of time points = 250 for rs-fMRI images; repetition time time = 7.5 ms, echo time = 3.7 ms, slice number =180, field of view =240×240 mm^2^, acquisition matrix = 256×200, and flip angle =8° for T1-weighted images). Data preprocessing was performed using the standard pipeline of the DPARSF software (31,32), including discarding the first 10 volumes, slice timing, head motion realignment, brain segmentation, spatial normalization, temporal filtering (0.01-0.10 Hz), as well as regressing out white matter and cerebrospinal fluid signals. Global signal regression was not performed as it is still a controversial preprocessing option in rs-fMRI studies (33). Subjects with excessive head motion were excluded from the analysis, as determined by a mean framewise-displacement (FD) (34) > 0.2 mm.

### 2.3. Temporal variability of FC

After preprocessing, the mean time series were extracted from each of the 116 regions of interest (ROIs) in the Automated Anatomical Labeling (AAL) atlas (35), which was validated (36) and widely used in functional neuroimaging studies (37,38). The names of all the 116 ROIs were listed in **Supplementary Table S1**.

As shown in **Figure 1**, to characterize the temporal variability of FC, all the time series were segmented into *n* nonoverlapping time windows with a length of *l*. Within each time window, a 116 × 116 pairwise Pearson correlation matrix was calculated to represent the FC between each pair of ROIs within that window. The temporal variability of regional FC architecture in each ROI could then be estimated by computing the mean values of its dissimilarities among different windows. Briefly, temporal variability of the regional FC architecture in ROI *k* is defined by Equation (1):

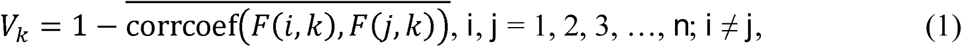

where n is the number of time windows, and *F*(*i, k*) is the vector characterizing the FC architecture between ROI *k* and the whole brain within the *i*th time window (**Figure 1**) (18,39,40).

**Figure 1.**
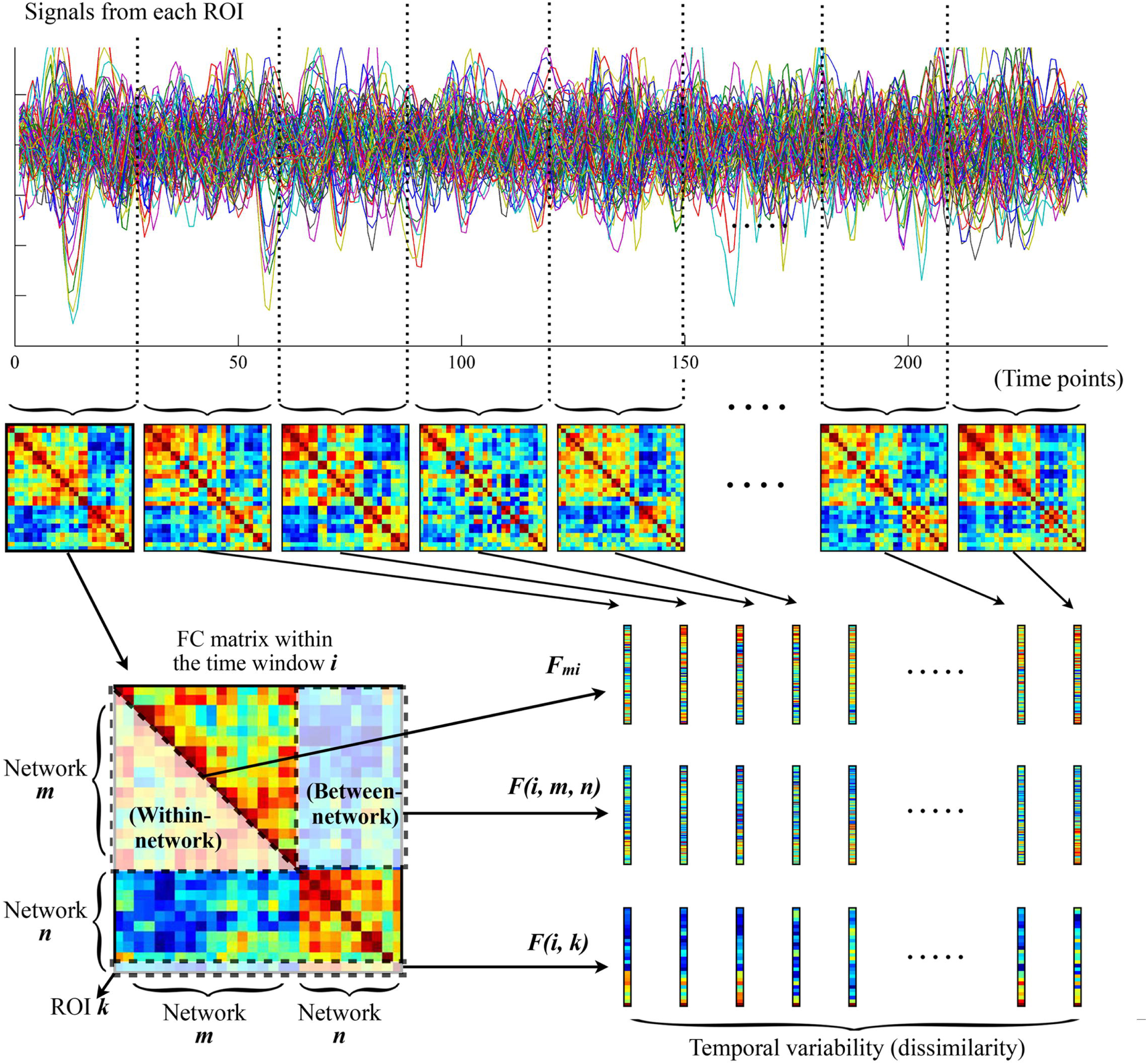
The procedures for computing temporal variabilities of FC patterns. Refer to the section 2.3, “Temporal Variability of FC” for details. FC, functional connectivity; ROI, region of interest.

The temporal variability of FC was further estimated at the network level following recently published procedures (22,23,41). First, all brain ROIs were assigned into eleven prior networks as defined in previous studies (42,43), including the sensorimotor network, visual network, auditory network, default-mode network, frontoparietal network, cingulo-opercular network, salience network, attention network, subcortical network, thalamus, and cerebellum (see **Supplementary Table S1** for details about the network assignments). Note that the thalamus and cerebellum were treated as two independent networks here, given that they were poorly defined into different networks as well as their special roles in the pathophysiologic mechanisms of psychotic disorders (22,44,45). The temporal variabilities of intra-network and inter-network FC architectures were then calculated among the above eleven networks. Similar with the regional FC variability for each ROI, the intra-network FC variability for a network *m* is defined by Equation (2):

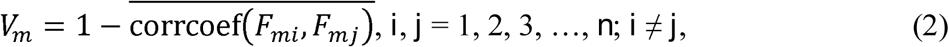

where *n* is the number of time windows, and *F*_*mi*_ is the vector characterizing the FC architecture between all ROIs belonging to the network *m* within the *i*th time window (**Figure 1**); the inter-network variability of FC between two networks *m* and *n* is defined by Equation (3):

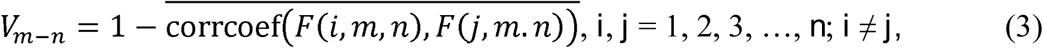

where *n* is the number of time windows, and *F*(*i, m, n*) is the vector characterizing the FC architecture between the networks *m* and *n* within the ith time window (**Figure 1**) (22,23,41).

To reduce the influences from window length and segmentation scheme, all the above temporal variabilities were calculated with a set of different window lengths (*l* = 21, 22, …, 30 volumes, equal to 42, 44, …, 60 seconds) and for each window length of *l*, we performed the segmentation for (*l-*1) times (starting from the first, second, …, (*l-*1)th volumes at each time). The final values of temporal variabilities were obtained by averaging all of these values. Note that such a selection of window lengths has been used in previous studies, and was suggested to be optimal for producing robust results (46,47). As the result, in each subject, we finally obtained the temporal variabilities of regional FC for each of the 116 ROIs, intra-network FC for each of the 11 networks, and inter-network FCs for each possible pair of networks. All these values of temporal variabilities range from 0 to 2, and a higher value suggests a higher variability.

### 2.4. Statistics

The demographic and clinical characteristics as well as mean FD were compared between groups using the two-sample *t*-test, Chi-square test or analysis of variance. Differences were considered significant at *p* < 0.05.

The temporal variabilities of FC patterns were compared between groups at all the regional, intra-network, and inter-network levels. The group differences were determined the by following statistic steps (46): 1) the analysis of covariance (ANCOVA) covarying for age, sex, education and head motion (mean FD) was firstly applied to detect the significant main effect; 2) post-hoc pairwise comparisons were adopted between all possible pairs of groups when the main effect was significant (*p* < 0.05); 3) the Bonferroni correction was applied to control the false-positive rate for multiple tests within the ANCOVA, and the groups differences were considered significant at corrected-*p* < 0.05.

For all the detected significant between-group differences, we further explored their possible relationships with the clinical and cognitive variables using Spearman’s rank correlation coefficient. Here, they were correlated with the illness duration, chlorpromazine equivalence, SAPS scores, SANS scores, HAMD scores, YMRS scores, WAIS-I scores and WAIS-DS scores in each group separately. The correlations were considered significant at *p* < 0.05.

## 3. Results

### 3.1. Demographic, clinical and head motion characteristics

As shown in **Table 1**, there were no significant differences among the three groups in age, sex and education (all *p* > 0.05). Shorter illness durations but higher antipsychotic doses (both *p* < 0.001) were observed in the schizophrenia patients compared with the bipolar disorder patients. Both the schizophrenia and bipolar disorder groups showed significantly lower WAIS-I and WAIS-DS scores (all *p* < 0.05, LSD post-hoc comparisons) compared with healthy controls, while there was no significant difference between the schizophrenia and bipolar disorder patients in WAIS-I and WAIS-DS scores. There was no significant difference among the three groups in head motion as measured by mean FD (*F* = 2.066, *p* = 0.130).

**Table 1.** Demographic, clinical and head motion characteristics of the three groups.

|  | Schizophrenia (*n* = 66) | Bipolar disorder (*n* = 53) | Healthy controls (*n* = 66) | Group comparisons |
| --- | --- | --- | --- | --- |
|  | (Mean ± SD) | (Mean ± SD) | (Mean ± SD) |  |
| Age (years) | 24.318 ± 6.127 | 25.340 ± 4.095 | 23.379 ± 4.416 | $F = 2.249, p = 0.108$ |
| Sex (male/female) | 38/28 | 26/27 | 28/38 | $\chi^2 = 3.044, p = 0.218$ |
| Education (years) | 13.152 ± 2.061 | 13.576 ± 2.578 | 14.076 ± 2.200 | $F = 2.745, p = 0.067$ |
| Illness duration (months) | 22.214 ± 24.972<sup>a</sup> | 56.987 ± 53.907 | / | $t = -4.281, p < 0.001$ |
| Antipsychotics (taking/not taking) | 63/3 | 38/15 | / | $\chi^2 = 3.609, p < 0.001$ |
| Fluoxetine equivalents (mg/day) | 228.762 ± 155.296 | 108.047 ± 119.101 | / | $t = 4.663, p < 0.001$ |
| SAPS scores | 18.231 ± 13.828 | / | / | / |
| SANS scores | 31.636 ± 27.810 | / | / | / |
| 17-item HAMD scores | / | 12.660 ± 9.265 | / | / |
| YMRS scores | / | 5.113 ± 8.257 | / | / |
| WAIS-I scores | 18.174 ± 4.143 | 19.123 ± 4.410 | 20.985 ± 4.774 | $F = 6.773, p < 0.001^b$ |
| WAIS-DS scores | 65.182 ± 15.493 | 70.321 ± 14.997 | 88.924 ± 13.014 | $F = 48.263, p < 0.001^b$ |
| Mean FD | 0.095 ± 0.038 | 0.082 ± 0.035 | 0.086 ± 0.032 | $F = 2.066, p = 0.130$ |

<sup>1</sup> The information on illness duration was available for 56 schizophrenia patients. <sup>2</sup> The LSD post-hoc comparisons set at $p < 0.05$ showed that schizophrenia < healthy controls, and bipolar disorder < healthy controls, while there was no significant difference between the schizophrenia and bipolar disorder groups. SD, standard deviation; SAPS, Scale for Assessment of Positive Symptoms; SANS, Scale for Assessment of Negative Symptoms; HAMD, Hamilton Rating Scale for Depression; YMRS, Young Mania Rating Scale; WAIS-I, the Information subtest of the Wechsler Adult Intelligence Scale; WAIS-DS, the Digit Symbol subtest of the Wechsler Adult Intelligence Scale; FD, framewise-displacement.

### 3.2. Differences in temporal variability of regional FC

As shown in **Supplementary Table S2** and **Figure 2**, for temporal variability of the regional FC, both the schizophrenia and bipolar disorder patients showed significantly higher variabilities in a number of subcortical ROIs, including the thalamus and regions of the basal ganglia (putamen/pallidum) compared with healthy controls; the schizophrenia patients additionally showed significantly higher variabilities for a number of ROIs located in the sensorimotor (precentral gyrus and postcentral gyrus), attention (inferior parietal lobule) and limbic (hippocampus and amygdala) areas than healthy controls, as well as a significantly lower variability in the superior frontal gyrus (medial orbital) than healthy controls and a significantly lower variability in the posterior cingulate gyrus than bipolar disorder patients (all corrected-*p* < 0.05).

**Figure 2.**
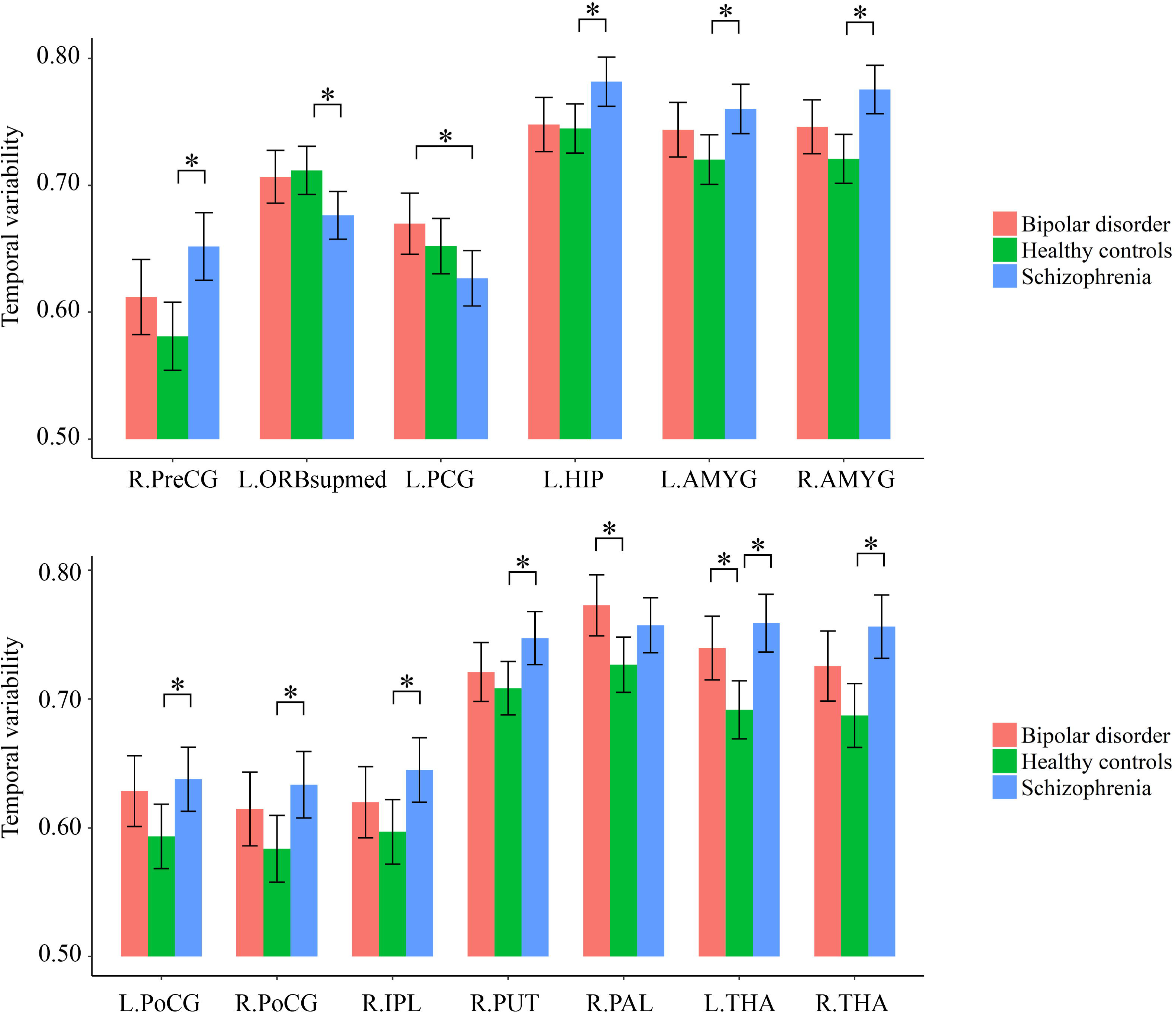
Group differences in the temporal variabilities of regional FC patterns. The error bars present the 95% confidence intervals, and the marks “*” indicate significant between-group differences with corrected p < 0.05. AMYG, amygdala; FC, functional connectivity; HIP, hippocampus; IPL, inferior parietal lobule; L, left hemisphere; ORBsupmed, superior frontal gyrus (medial orbital); PAL, pallidum; PCG, posterior cingulate gyrus; PoCG, postcentral gyrus; PreCG, precentral gyrus; PUT, putamen; R, right hemisphere; THA, thalamus.

### 3.3. Differences in temporal variability of intra- and inter-network FC

As shown in **Supplementary Table S3** and **Figure 3**, for temporal variabilities of the intra-network FC within particular networks and inter-network FC between particular pairs of networks, both the schizophrenia and bipolar disorder patients showed a significantly higher variability for inter-network FC between the sensorimotor network and thalamus compared with healthy controls; the schizophrenia patients additionally showed significantly higher variabilities of both intra-network and inter-network FC than healthy controls for several networks and pairs of networks, which mainly involved the sensorimotor, visual and subcortical (including the thalamus) networks (all corrected p < 0.05).

**Figure 3.**
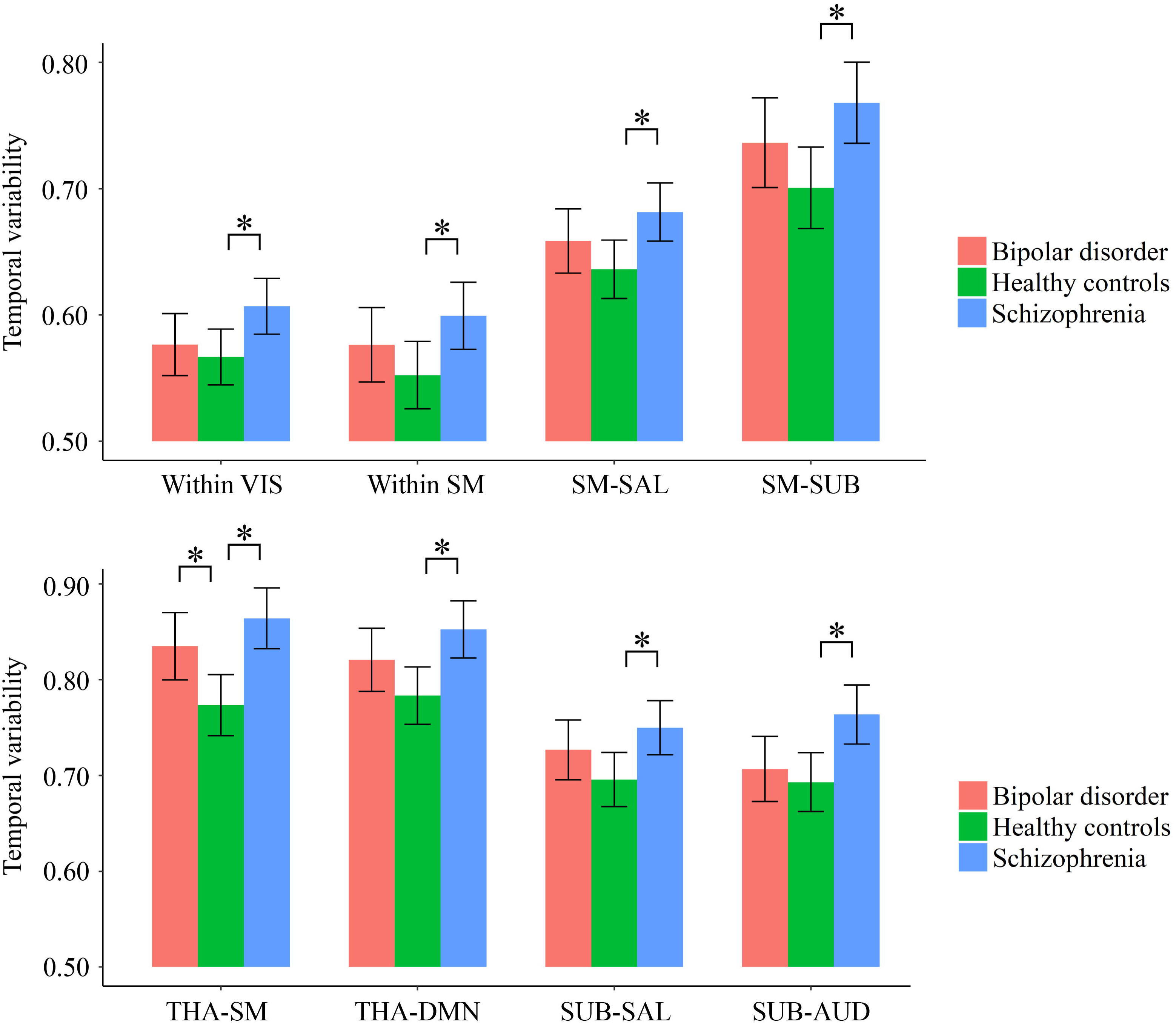
Group differences in the temporal variabilities of intra-network and inter-network FC patterns. The error bars present the 95% confidence intervals, and the marks “*” indicate significant between-group differences with corrected p < 0.05. AUD, auditory network; DMN, default mode network; FC, functional connectivity; SAL, salience network; SM, sensorimotor network; SUB, subcortical network; THA, thalamus; VIS, visual network.

### 3.4. Correlations

As shown in **Figures 4A** and **4B**, in the group of schizophrenia patients, significant correlations were found between temporal variability of regional FC for left hippocampus and the SANS scores (Spearman’s rho = 0.330, *p* = 0.007, uncorrected for multiple tests), as well as between temporal variability of the inter-network FC between subcortical and auditory networks and the WAIS-I scores (Spearman’s rho = −0.286, *p* = 0.020, uncorrected for multiple tests). No significant correlations were found in the groups of healthy controls and bipolar disorder patients (*p* > 0.05).

**Figure 4.**
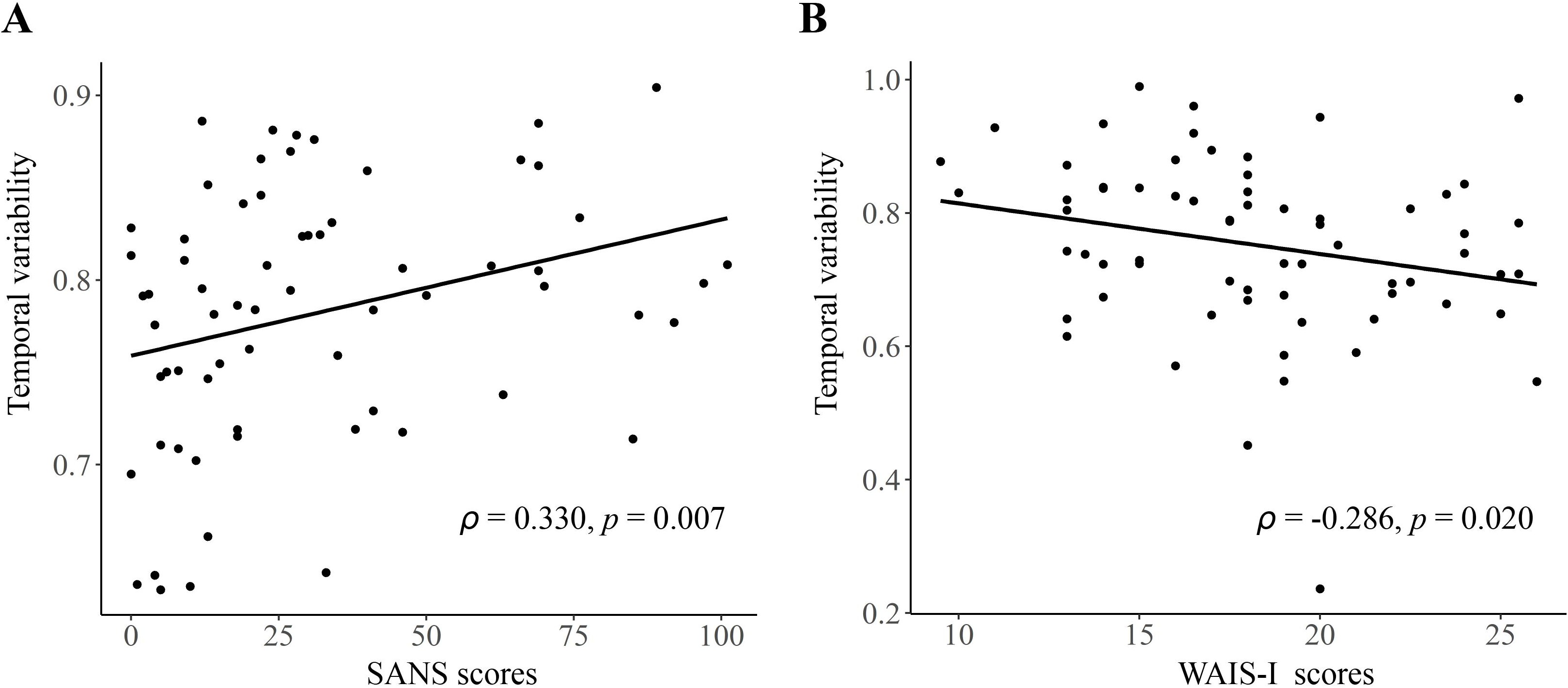
The detected significant correlations in schizophrenia patients. **(A)** Correlation between the temporal variability of regional FC for left hippocampus and the Scale for Assessment of Negative Symptoms (SANS) scores. **(B)** Correlation between the temporal variability of inter-network FC between subcortical and auditory networks and the Information Subtest of the Wechsler Adult Intelligence Scale (WAIS-I) scores. The Spearman’s correlation coefficients (ρ) and p values are presented on figures.

## 4. Discussion

In this study, we explored the common and specific changes in dynamic local and large-scale resting-state FC, as characterized by altered temporal variabilities, across the schizophrenia and bipolar disorder. Our results provide some innovative findings on the dynamic functional architecture of the brain for these two severe mental disorders: firstly, we found that both the schizophrenia and bipolar disorder patients showed increased regional FC variabilities in a number of subcortical areas involving the thalamus and regions of basal ganglia, as well as increased inter-network FC variability between the sensorimotor cortices and thalamus; secondly, some specific abnormalities were found to present only in the schizophrenia group, at both regional and network levels in a wider range. These findings provide valuable information for improving our insight into the neuropathology of these disorders from a dynamic brain functional perspective.

Our first important finding is that both the schizophrenia and bipolar disorder patients exhibited similar increased temporal variabilities of local FC in the thalamus (**Figure 2**), as well as of inter-network FC between the thalamus and sensorimotor cortices (**Figure 3**). It is noteworthy that shared neural disturbances in thalamo-cortical communications across schizophrenia and bipolar disorder, as characterized by similar over-connectivity between the thalamus and sensorimotor regions, have been repeatedly reported in several previous conventional static rs-fMRI studies (3,4). Our results, therefore, may extend such findings to the context of dynamic resting-state FC for the first time to our knowledge. The thalamus is known as a “relay station” for almost all motor and sensory information flow from and to the cortex, where the information is further processed for high-order brain functions (3,48). Specifically, aberrant communications between the thalamus and sensorimotor network were presumed to reflect a sensory gating deficit which leads to abnormal sensory information flow through the thalamus to the cortex (4,45,49). The observed increased temporal variability of thalamo-sensorimotor connectivity could thus point to such a sensory gating deficit, as abnormally increased temporal variability of FC was suggested to reflect excessive fluctuations in brain activities and inappropriate processing of information (22). As notably reported in both the schizophrenia and bipolar disorder patients (50–52), the sensory gating deficit has been suggested to partly underlie the cognitive and perceptual symptoms in the disorders (3,53). Therefore, our dynamic FC findings may further support the hypothesis that thalamo-sensorimotor connectivity disturbances and sensory gating deficits are common neurobiological features shared by schizophrenia and bipolar disorder (4,50).

In the present study, we also found that both the schizophrenia and bipolar disorder patients showed increased local FC variability in regions of the basal ganglia (putamen and pallidum) (**Figure 2**). The basal ganglia is a group of subcortical nuclei (putamen, pallidum, caudate nucleus, substantia nigra, and subthalamic nucleus) that involves a variety of brain functions such as motor control, learning, and execution (54). The functional and structural abnormalities of basal ganglia have been widely reported to be associated with psychotic symptoms such as delusions in schizophrenia patients (55–57), and also present in psychotic bipolar disorder patients (58). Therefore, our findings of such shared alterations in the basal ganglia may be reflective of common functional deficits in both the schizophrenia and bipolar disorder. These findings, together with the observed shared alterations in the thalamo-sensorimotor circuit, may partly help to explain the overlap clinical features in these two disorders.

Besides the above shared alterations in both patient groups, some specific alterations in a much wider range were found to present in only the schizophrenia patients. These include widespread increased FC variabilities at both regional and network levels, involving the sensorimotor, visual, attention, limbic and subcortical areas, as well as decreased regional FC variability in a number of areas comprising the default-mode network such as posterior cingulate gyrus and superior frontal gyrus (medial orbital part) (**Figure 2** and **Figure 3**). Generally, these results are highly consistent with the findings from another recent study (22), which reported that schizophrenia patients had significantly increased FC variabilities in sensory and perceptual systems (including the sensorimotor network, visual network, attention network, and thalamus) and decreased FC variabilities in high-order networks (including the default-mode and frontal–parietal networks) than healthy subjects at both regional and network levels. Moreover, these alterations were found to be related to patients’ clinical symptoms and cognitive deficits both in the present study (**Figure 4**) and prior research (22). Therefore, our results further support the recent opinion that such widespread aberrant dynamic brain network reconfigurations may constitute a potential reliable biomarker for schizophrenia, suggestive of impaired abilities in processing inputs in sensory/perceptual systems and integrating information in high-order networks, which may underlie the perceptual and cognitive deficits in schizophrenia (22,59). As for the bipolar disorder patients in the present study, FC variabilities in these regions and networks did not differ significantly from either of the other groups, which fell in the intermediate range between those of healthy controls and schizophrenia patients (**Figure 2** and **Figure 3**). Thus, we propose that such findings may offer support for the hypothesis of a psychosis continuum between schizophrenia and bipolar disorder, with more severe brain deficits and disabling symptoms in schizophrenia compared to bipolar disorder (60,61). However, future investigation with a larger sample size and a higher statistical power is required to confirm if these changes would be significant in patients with bipolar disorder, as compared to healthy controls and schizophrenia patients.

There are several limitations of the present study and future research directions which should be noted. First, as mentioned before, our sample size is relatively small and the results should be further verified in future work with a larger sample to increase the reliability and statistical power (62). Second, the illness duration and doses of antipsychotics were not matched between the schizophrenia and bipolar disorder groups. Although no significant correlations were found between them and any detected group differences, which suggests that the observed group differences are unlikely to be mainly driven by medications or long-term hospitalizations, further studies using drug-naïve or matched samples for medication and illness duration are warranted to exclude their possible effects. Third, a number of previous studies have pointed out that the psychotic bipolar disorder may be a special phenotype from non-psychotic bipolar disorder (63,64). In the current sample, the records of psychotic symptom histories are unavailable for most bipolar disorder patients. Future studies are necessary to replicate our results and to compare between psychotic and non-psychotic bipolar disorder patients. Fourth, while many important results were found in the thalamus, we examined the thalamus as a single entity by the AAL atlas. However, the thalamus can be anatomically subdivided into multiple distinct nuclei with different FC patterns (44,65). Future studies to investigate the temporal variability of thalamo-cortical FC patterns within different sub-regions in the thalamus would further improve our understanding of its important role in the schizophrenia and bipolar disorder.

In conclusion, we explored the common and specific changes in dynamic features of FC, as characterized by temporal variabilities of FC patterns involved in specific brain regions or large-scale brain networks, in schizophrenia and bipolar disorder patients. We found that both the schizophrenia and bipolar disorder patients showed significantly increased regional FC variabilities in subcortical areas including the thalamus and basal ganglia, as well as increased inter-network FC variability between the sensorimotor cortices and thalamus. More widespread significant alterations were found to present in only the schizophrenia group, including increased FC variabilities in the sensorimotor, visual, attention, limbic and subcortical areas at both regional and network levels, as well as decreased regional FC variability in the default-mode areas. The observed alterations shared by schizophrenia and bipolar disorder may help to explain their overlap clinical features; meanwhile, the schizophrenia-specific abnormalities in a wider range could potentially support the hypothesis of a psychosis continuum between schizophrenia and bipolar disorder, that schizophrenia is associated with more severe functional brain deficits compared to bipolar disorder.

## AUTHOR CONTRIBUTIONS

Authors YL and WP designed the study and carried out the analysis. YL, ZL, GW, ZX, YP, XC, XH and WP contributed to the data collection. YL wrote the first draft of manuscript. ZL, CKC and WP contributed to the final manuscript. All authors have read and agreed to the published version of the manuscript.

## ACKNOWLEDGMENTS

We thank Zhong He (Department of Radiology of Second Xiangya Hospital, Central South University) for his assistance in rs-fMRI data acquisition.

## SUPPLEMENTARY MATERIAL

The Supplementary Material for this article can be found online.

