## Supplementary Material for "Altered Temporal Variability of Local and Large-scale Resting-state Brain Functional Connectivity Patterns in Schizophrenia and Bipolar Disorder"

**Supplementary Table S1.** List of the 116 regions of interest used in the present study and their network affiliations based on prior research. Odd and even numbers represent left and right hemispheres, respectively.

| Index | Corresponding Brain Region | Network affiliation |
| --- | --- | --- |
| (1,2) | Precentral gyrus | Sensorimotor |
| (3,4) | Superior frontal gyrus, dorsolateral | Frontoparietal |
| (5,6) | Superior frontal gyrus, orbital part | Frontoparietal |
| (7,8) | Middle frontal gyrus | Salience/frontoparietal/attention |
| (9,10) | Middle frontal gyrus, orbital part | Frontoparietal |
| (11,12) | Inferior frontal gyrus, opercular part | Cingulo-opercular |
| (13,14) | Inferior frontal gyrus, triangular part | Salience/frontoparietal/attention |
| (15,16) | Inferior frontal gyrus, orbital part | None |
| (17,18) | Rolandic operculum | Auditory/cingulo-opercular |
| (19,20) | Supplementary motor area | Sensorimotor |
| (21,22) | Olfactory cortex | None |
| (23,24) | Superior frontal gyrus, medial | Default-mode |
| (25,26) | Superior frontal gyrus, medial orbital | Default-mode |
| (27,28) | Gyrus rectus | None |
| (29,30) | Insula | Salience/cingulo-opercular |
| (31,32) | Anterior cingulate and paracingulate gyri | Default-mode/salience |
| (33,34) | Median cingulate and paracingulate gyri | Salience/cingulo-opercular |
| (35,36) | Posterior cingulate gyrus | Default-mode |
| (37,38) | Hippocampus | None |
| (39,40) | Parahippocampal gyrus | Default-mode |
| (41,42) | Amygdala | None |
| (43,44) | Calcarine fissure and surrounding cortex | Visual |
| (45,46) | Cuneus | Visual |
| (47,48) | Lingual gyrus | Visual |
| (49,50) | Superior occipital gyrus | Visual |
| (51,52) | Middle occipital gyrus | Visual |
| (53,54) | Inferior occipital gyrus | Visual |
| (55,56) | Fusiform gyrus | Visual |
| (57,58) | Postcentral gyrus | Sensorimotor |
| (59,60) | Superior parietal gyrus | Salience/attention |
| (61,62) | Inferior parietal, but supramarginal and angular gyri | Frontoparietal/attention |
| (63,64) | Supramarginal gyrus | Auditory/cingulo-opercular |
| (65,66) | Angular gyrus | Default-mode |
| (67,68) | Precuneus | Default-mode |
| (69,70) | Paracentral lobule | Sensorimotor |
| (71,72) | Caudate nucleus | Subcortical |
| (73,74) | Lenticular nucleus, putamen | Subcortical |
| (75,76) | Lenticular nucleus, pallidum | Subcortical |
| (77,78) | Thalamus | Thalamus |
| (79,80) | Heschl gyrus | Auditory |
| (81,82) | Superior temporal gyrus | Auditory/attention |
| (83,84) | Temporal pole: superior temporal gyrus | Cingulo-opercular |
| (85,86) | Middle temporal gyrus | Default-mode |
| (87,88) | Temporal pole: middle temporal gyrus | Default-mode |
| (89,90) | Inferior temporal gyrus | None |
| (91,92) | Cerebellum crus | Cerebellum |
| (93,94) | Cerebellum crus | Cerebellum |
| (95,96) | Cerebellum | Cerebellum |
| (97,98) | Cerebellum | Cerebellum |
| (99,100) | Cerebellum | Cerebellum |
| (101,102) | Cerebellum | Cerebellum |
| (103,104) | Cerebellum | Cerebellum |
| (105,106) | Cerebellum | Cerebellum |
| (107,108) | Cerebellum | Cerebellum |
| (109,110) | Vermis | Cerebellum |
| (111,112) | Vermis | Cerebellum |
| (113,114) | Vermis | Cerebellum |
| (115,116) | Vermis | Cerebellum |

**Supplementary Table S2.** The detected significant between-group differences in temporal variabilities of regional functional connectivity for particular regions of interest.

| Region of interest | Main effect of group | Significant post-hoc pairwise comparisons^1^ |
| --- | --- | --- |
| Right precentral gyrus | *F* = 6.625, *p* = 0.002 | Schizophrenia > healthy controls (*p* = 0.001) |
| Left superior frontal gyrus, medial orbital | *F* = 3.865, *p* = 0.023 | Schizophrenia < healthy controls (*p* = 0.032) |
| Left posterior cingulate gyrus | *F* = 3.482, *p* = 0.033 | Schizophrenia < bipolar disorder (*p* = 0.030) |
| Left hippocampus | *F* = 4.137, *p* = 0.018 | Schizophrenia > healthy controls (*p* = 0.029) |
| Left amygdala | *F* = 3.979, *p* = 0.020 | Schizophrenia > healthy controls (*p* = 0.017) |
| Right amygdala | *F* = 7.617, *p* = 0.001 | Schizophrenia > healthy controls (*p* = 0.0004) |
| Left postcentral gyrus | *F* = 3.277, *p* = 0.040 | Schizophrenia > healthy controls (*p* = 0.047) |
| Right postcentral gyrus | *F* = 3.547, *p* = 0.031 | Schizophrenia > healthy controls (*p* = 0.027) |
| Right inferior parietal lobule | *F* = 3.472, *p* = 0.033 | Schizophrenia > healthy controls (*p* = 0.028) |
| Right putamen | *F* = 3.483, *p* = 0.033 | Schizophrenia > healthy controls (*p* = 0.031) |
| Right pallidum | *F* = 4.246, *p* = 0.016 | Bipolar disorder > healthy controls (*p* = 0.015) |
| Left thalamus | *F* = 8.915, *p* = 0.0002 | Schizophrenia > healthy controls (*p* = 0.0002), bipolar disorder > healthy controls (*p* = 0.017) |
| Right thalamus | *F* = 7.386, *p* = 0.001 | Schizophrenia > healthy controls (*p* = 0.001) |

^1^ The *p* values were Bonferroni-corrected for multiple tests within the analysis of covariance.

**Supplementary Table S3.** The detected significant between-group differences in temporal variabilities of intra-network and inter-network functional connectivity for particular networks or pairs of networks.

| Network/pair of networks | Main effect of group | Significant post-hoc pairwise comparisons^1^ |
| --- | --- | --- |
| Visual | *F* = 3.359, *p* = 0.037 | Schizophrenia > healthy controls (*p* = 0.041) |
| Sensorimotor | *F* = 3.083, *p* = 0.048 | Schizophrenia > healthy controls (*p* = 0.049) |
| Sensorimotor-salience | *F* = 3.642, *p* = 0.028 | Schizophrenia > healthy controls (*p* = 0.023) |
| Sensorimotor-subcortical | *F* = 4.134, *p* = 0.018 | Schizophrenia > healthy controls (*p* = 0.014) |
| Thalamus-sensorimotor | *F* = 7.907, *p* = 0.001 | Schizophrenia > healthy controls (*p* = 0.0004), bipolar disorder > healthy controls (*p* = 0.036) |
| Thalamus-default-mode | *F* = 5.060, *p* = 0.007 | Schizophrenia > healthy controls (*p* = 0.005) |
| Subcortical-salience | *F* = 3.464, *p* = 0.033 | Schizophrenia > healthy controls (*p* = 0.028) |
| Subcortical-auditory | *F* = 5.529, *p* = 0.005 | Schizophrenia > healthy controls (*p* = 0.006) |

^1^ The *p* values were Bonferroni-corrected for multiple tests within the analysis of covariance.
